# Amplification of olfactory transduction currents implements sparse stimulus encoding

**DOI:** 10.1101/2024.10.11.617893

**Authors:** Kai Clane Belonio, Eyerusalem S. Haile, Zach Fyke, Lindsay Vivona, Vaibhav Konanur, Joseph D. Zak

## Abstract

Sensory systems must perform the dual and opposing tasks of being sensitive to weak stimuli while also maintaining information content in dense and variable sensory landscapes. This occurs in the olfactory system, where OSNs are highly sensitive to low concentrations of odors and maintain discriminability in complex odor environments. How olfactory sensory neurons (OSNs) maintain both sensitivity and sparsity is not well understood. Here, we investigated whether the calcium-activated chloride channel, TMEM16B, may support these dual roles in OSNs. We used multiphoton microscopy to image the stimulus-response density of OSNs in the olfactory epithelium. In TMEM16B knockout mice, we found that sensory representations were denser, and the magnitude of OSN responses was increased. Behaviorally, these changes in sensory representations were associated with an increased aversion to the odorant trimethylamine, which switches perceptual valence as its concentration increases, and a decreased efficiency of olfactory-guided navigation. Together, our results indicate that the calcium-activated chloride channel TMEM16B sparsens sensory representations in the peripheral olfactory system and contributes to efficient integrative olfactory-guided behaviors.

## Introduction

Each sensory modality is organized to detect minute quantities of environmental stimuli. To achieve such sensitivity, sensory transduction is amplified through biochemical and/or electrochemical signaling cascades (Kleene, 2008; Yildiz and Khanna, 2012; Palczewski, 2014). Olfactory sensory neurons (OSNs) use both types of cascades to detect odorants at exceedingly low concentrations (Kleene, 2008), but olfactory sensation must also operate in higher concentration regimes. How signaling cascades are tuned to maintain efficient coding in variable concentration environments is not well understood. For example, a given odorant can interact with a range of olfactory receptors (ORs) across a spectrum of binding affinities, thereby generating dense OSN ensemble activity. However, efficient information transfer demands readout from a sparse subset of well-activated OSNs (Davison and Katz, 2007; Betkiewicz et al., 2020; Zavatone-Veth et al., 2023). If transduction current amplification functions to drive up population activity at the sensory periphery as concentrations of odorants increase, how is sparsity maintained?

In the olfactory system, volatilized odorant molecules bind with G-protein coupled ORs found on the cilia of OSNs within the nasal mucosa (Mombaerts, 2004). This causes a canonical signaling cascade leading to the opening of cyclic nucleotide-gated channels, allowing for the influx of Na^+^ and Ca^2+^ ions, the initial components of the olfactory transduction current (Nakamura and Gold, 1987; Bradley et al., 2005). The initial influx of cations results in only a modest OSN membrane depolarization (Hengl et al., 2010; Billig et al., 2011; Neureither et al., 2017). Then, as a final step, accumulating Ca^2+^ ions engage the second component of the electrochemical cascade, the calcium-activated chloride channel, TMEM16B (also called anoctamin-2; Kleene and Gesteland, 1991). In OSNs, the Cl^-^ gradient favors the intracellular compartment (Kaneko and H., 2004), resulting in the efflux of negative charge and amplifying the initial transduction current (**Figure 1**). Given this behavior, TMEM16B was thought to primarily play a role as a signal amplifier.

**Figure 1.**
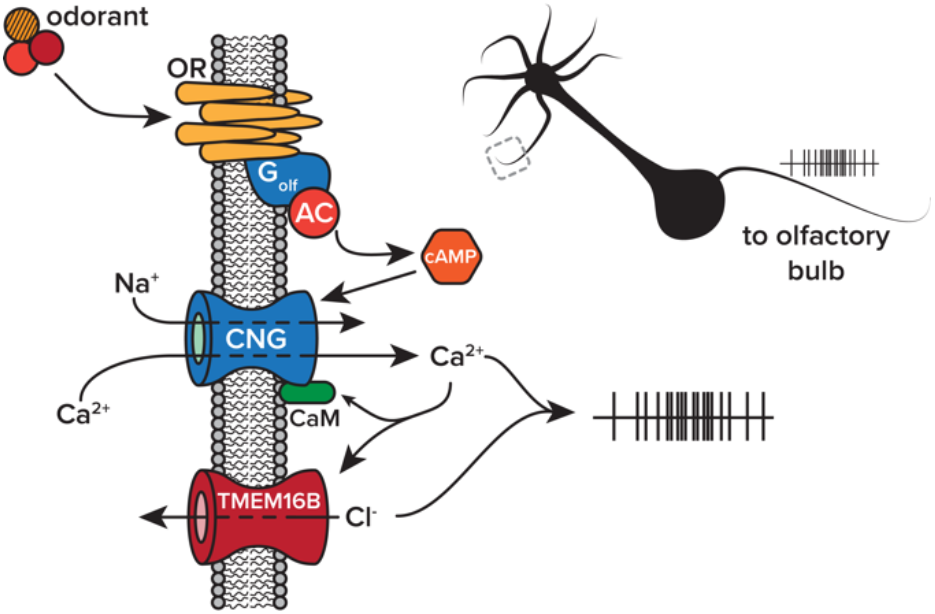
Olfactory transduction cascade. Volatilized odorant molecules activate a biochemical and electrochemical transduction cascade in olfactory sensory neurons. The final step of the cascade, efflux of chloride through TMEM16B, amplifies transduction currents through cyclic nucleotide gate (CNG) channels. Abbreviations, olfactory receptor (OR), adynyl cyclase III (AC), cyclic adenosine monophosphate (cAMP), calmodulin (CaM).

It was initially thought that current amplification through TMEM16B ensured signaling fidelity from the periphery to central circuits. However, evidence has recently emerged that TMEM16B paradoxically limits action potential generation in OSNs despite amplifying transduction currents (Pietra et al., 2016; Zak et al., 2018; Reisert et al., 2024). We hypothesized that TMEM16B operates as a shunt on OSN output, thereby functioning to sparsen dense peripheral inputs to the olfactory system. Here, we studied the role that TMEM16B plays in encoding sensory stimuli by measuring OSN activity *in vivo* using multiphoton microscopy and testing the role of TMEM16B in naturalistic odor-guided behaviors.

## Results

### Functional imaging of OSN output in the olfactory epithelium of live mice

How does transduction current amplification contribute to sensory neuron activity in live animals at the level of individual neurons? Previous characterizations of TMEM16B’s contribution to olfactory transduction relied on somatic recordings of dissociated cells or population measurements from OSN axon terminals in the olfactory bulb (Stephan et al., 2009; Pietra et al., 2016; Zak et al., 2018; Reisert et al., 2024). Under both conditions, the transmembrane chloride gradient is unlikely to be consistent with the mucosal surroundings of cilia in the olfactory epithelium. This could confound the observation that TMEM16B functions to limit OSN spike generation (Pietra et al., 2016; Zak et al., 2018; Reisert et al., 2024). We measured odorant-evoked response at the somata of individual OSNs in the olfactory epithelium of live mice using multiphoton microscopy. In mice heterozygous for the *OMP-GCaMP3* allele (Isogai et al., 2011) and either *Tmem16b*^*+/+*^ or *Tmem16b*^*-/-*^, we used a bone thinning procedure to gain optical access to the olfactory epithelium (Zak et al., 2020, 2024; Zak, 2022). We then individually delivered a panel of 32 unique monomolecular odorants to each cohort (**Figure 2A-C**). We designed our odorant panel to generate a range of OSN response densities.

**Figure 2.**
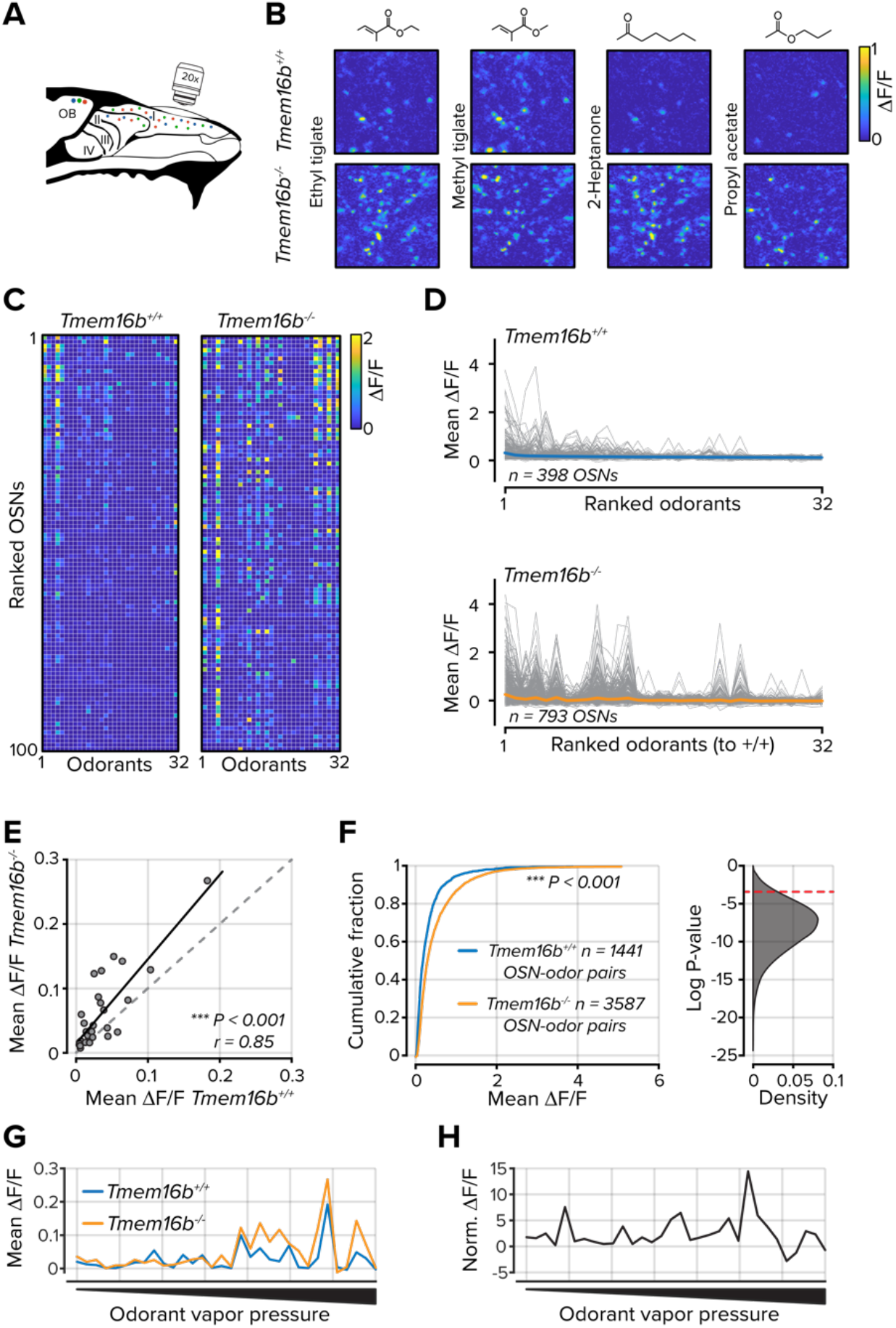
TMEM16B limits OSN activity in the olfactory epithelium. **A**. Schematic of olfactory epithelium imaging. **B**. Example ΔF/F images from the olfactory epithelium in *Tmem16b*^*+/+*^ and *Tmem16b*^*-/-*^ mice for four odorants. **C**. Mean ΔF/F signals for each of 32 odorants in 100 randomly selected OSNs. Data are ranked to the mean of all odorants.**D**. Tuning curves for each OSN (gray lines). The mean of all OSNs is shown as the colored lines. **E**. Scatter plot of mean ΔF/F values for each of 32 odorants. **F**. *Left*, the cumulative distribution of all OSN-odorant pairs. *Right*, distribution of 50,000 bootstrapped *P* values from the distributions at left. The dashed red line indicates a *P* value of 0.05. **G**. Mean responses for each odorant rank-ordered to odorant vapor pressure. **H**. *Tmem16b*^*-/-*^ OSN responses normalized to *Tmem16b*^*+/+*^ OSNs and ranked by odorant vapor pressure.

In 19 out of 32 odorants, OSNs in *Tmem16b*^*-/-*^ mice had larger responses (Bonferroni corrected *P* values < 0.0016 with rank sum test; **Figure 2D,E**). Considering all OSN-odorant pairs, OSNs in *Tmem16b*^*-/-*^ mice responded to odorants with larger Ca^2+^ transients across all OSN-odorant pairs (*n* = 1345 *Tmem16b*^*+/+*^ and 3051 *Tmem16b*^*-/-*^ OSN-odorant pairs; *P* < 0.001; Kolmogorov-Smirnov test; **Figure 2F**).

TMEM16B may have differential effects depending on the intensity of a stimulus. For instance, at low odorant concentrations, the channel may act as an output amplifier, while at higher concentrations, it may limit OSN activity. When generating our odorant panel, we used a nominal 1% v/v dilution for each. However, the panel consisted of odorants with vapor pressures that spanned greater than six orders of magnitude. Therefore, we could compare OSN responses across a range of effective odorant concentrations. We found a strong relationship between odorant vapor pressure and the mean OSN population activity in both genotypes (**Figure 2G**). However, when we normalized *Tmem16b*^*-/-*^ OSN activity to *Tmem16b*^*+/+*^ OSNs, we did not find weaker OSN responses in *Tmem16b*^*-/-*^ OSNs at the lowest vapor pressures and effective concentrations (**Figure 2H**). Among the 16 odorants with the lowest vapor pressures, *Tmem16b*^*-/-*^ OSNs had stronger responses for each, indicating that TMEM16B does not amplify OSN output at low odorant concentrations. Our data are consistent with previous evidence that TMEM16B functions as a shunt for OSN output despite amplifying OSN transduction currents (Pietra et al., 2016; Zak et al., 2018; Reisert et al., 2024). However, it is not known how the increased output from OSNs affects their tuning properties and stimulus representations.

### Stimulus representations are reconfigured in the absence of TMEM16B

Next, we considered how enhanced activity modifies OSN tuning and stimulus encoding. Using the same panel of 32 odorants, we first measured lifetime sparseness from each OSN in both cohorts (**Figure 3A**). Lifetime sparseness provides a measure of the tuning breadth for individual OSNs. Despite their increased output, lifetime sparseness in *Tmem16b*^*-/-*^ OSNs was less than in *Tmem16b*^*+/+*^ OSNs (mean lifetime sparseness = 0.19 ± 0.006; *n* = 834 in *Tmem16b*^*+/+*^ OSNs and 0.16 ± 0.004; *n* = 1168 in *Tmem16b*^*-/-*^ OSNs; *P* < 0.001, Kolmogorov-Smirnov test; **Figure 3A**), indicating that *Tmem16b*^*-/-*^ OSNs are in fact less odorant selective.

**Figure 3.**
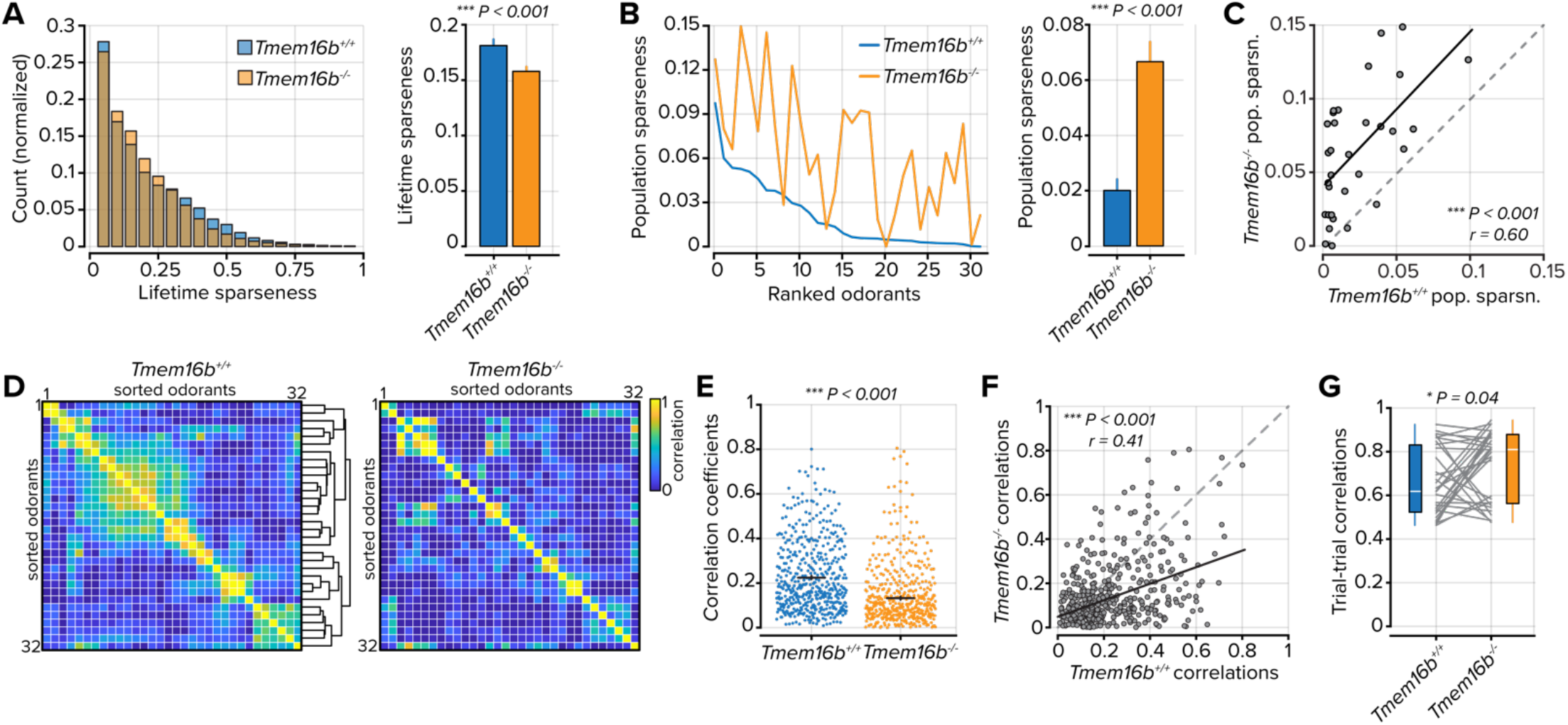
Odorant tuning and representations in OSNs. **A**. *Left*, distribution of lifetime sparseness values for each OSN. *Right*, average lifetime sparseness. The error bars represent the standard error of the mean. **B**. *Left*, population sparseness for each of 32 odorants rank-ordered to *Tmem16b*^*+/+*^ OSNs. Right, mean population sparseness. **C**. Scatter plot of the relationship between population sparseness for each odorant in *Tmem16b*^*+/+*^ OSNs and *Tmem16b*^*-/-*^ OSNs. **D**. Odorant-odorant correlations in *Tmem16b*^*+/+*^ OSNs and *Tmem16b*^*-/-*^ OSNs. White lines bound individual odorants, containing three to seven trials. Hierarchical clustering was used to group similar odorants in *Tmem16b*^*+/+*^ OSNs, and the clusters were then used to group datasets in *Tmem16b*^*-/-*^ OSNs. **E**. Odorant-odorant correlation coefficients from *part D*. Black lines represent the mean and the standard error of the mean. **F**. Scatter plot of the relationship between odorant-odorant correlations in *Tmem16b*^*+/+*^ and *Tmem16b*^*-/-*^ OSNs. **G**. Trial-to-trial correlation values for each of the 32 odorants. The white line represents the median.

Next, we used a second metric, population sparseness, to measure the density of odorant responses across the entire OSN population. Population sparseness measures not only the fraction of active units but also the degree of activation across all units. Here, we found that population sparseness was greater in *Tmem16b*^*-/-*^ OSNs (mean population sparseness = 0.02 ± 0.004 in *Tmem16b*^*+/+*^ OSNs and 0.06 ± 0.007 in *Tmem16b*^*-/-*^ OSNs; *n* = 32 odorants; *P* < 0.001, Sign-rank test; **Figure 3B**). Between *Tmem16b*^*+/+*^ *and Tmem16b*^*-/-*^ OSNs, there was also a strong relationship between population sparseness measures on a per odorant basis – odorants that generated dense population activity in *Tmem16b*^*+/+*^ OSNs were also found to generate dense population activity in *Tmem16b*^*-/-*^ OSNs (*r* = 0.60; *n* = 32 odorants; *P* < 0.001; **Figure 3C**).

The increased density of OSN responses may reconfigure representational similarity between odorants. To test this, we calculated pairwise odorant-odorant correlation coefficients in both cohorts of mice. We first computed the pairwise similarity between each of the 32 odorants in *Tmem16b*^*+/+*^ OSNs (**Figure 3D**, *left*). We then used hierarchical clustering to group odorants with similar OSN response profiles. When we imposed clusters from *Tmem16b*^*+/+*^ OSNs onto correlation coefficients measured in *Tmem16b*^*-/-*^ OSNs, the odorant relationships were largely unpreserved between groups (**Figure 3D**, *right*). We also found that in *Tmem16b*^*-/-*^ OSNs, there was a general decorrelation between odorant pairs (mean correlation coefficient = 0.23 ± 0.007 in *Tmem16b*^*+/+*^ OSNs and 0.14 ± 0.006 in *Tmem16b*^*-/-*^ OSNs; *n* = 496 odorant-odorant pairs; *P* < 0.001, Sign-rank test; **Figure 3E**). This difference was systematic as the odorant pair correlations strongly favored *Tmem16b*^*+/+*^ OSNs when compared between cohorts (*r* = 0.41; *P* < 0.001; **Figure 3F**).

As a final component of this analysis, we considered how TMEM16B may contribute to the stability of odorant representations across repeated sampling events. We delivered our odorant panel between three and seven times for each imaging field. We then measured the representational similarity of each odorant across repeats. On average, odorant representations were more stable in *Tmem16b*^*-/-*^ OSNs (mean trial-trial correlation = 0.65 ± 0.028 in *Tmem16b*^*+/+*^ OSNs and 0.73 ± 0.028 in *Tmem16b*^*-/-*^ OSNs; *n* = 32 odorants; *P* = 0.04, Sign-rank test; **Figure 3G**). Our results indicate that TMEM16B contributes to the density of OSN activity and odorant representations in the sensory periphery.

### Sustained output in Tmem16b^-/-^ OSNs

Our previous measures relied on the peak of the measured calcium signal, which is related to the firing rate of a neuron (Deneux et al., 2016; Huang et al., 2021). The sparsity of a sensory representation can also be measured by sustained activity. Next, we compared the temporal dynamics of *Tmem16b*^*+/+*^ and *Tmem16b*^*-/-*^ OSNs. From all OSNs that were modulated above baseline activity (see *Methods*), we measured the temporal profiles of their stimulus responses (**Figure 4A**). The average response of *Tmem16b*^*-/-*^ OSNs was larger than *Tmem16b*^*+/+*^ OSNs, and when normalized, their Ca^2+^ transients were longer lasting (peak normalized area 9.30 ± 0.15; *n* = 3820 in *Tmem16b*^*+/+*^ OSNs and 13.11 ± 0.13; *n* = 7621 in *Tmem16b*^*-/-*^ OSNs; *P* < 0.001; Kolmogorov-Smirnov test; **Figure 4B**). However, when using principal component analysis (PCA)to compare all OSN-odorant responses, we found little difference in the trajectory between the first and second components (**Figure 4C**).

**Figure 4.**
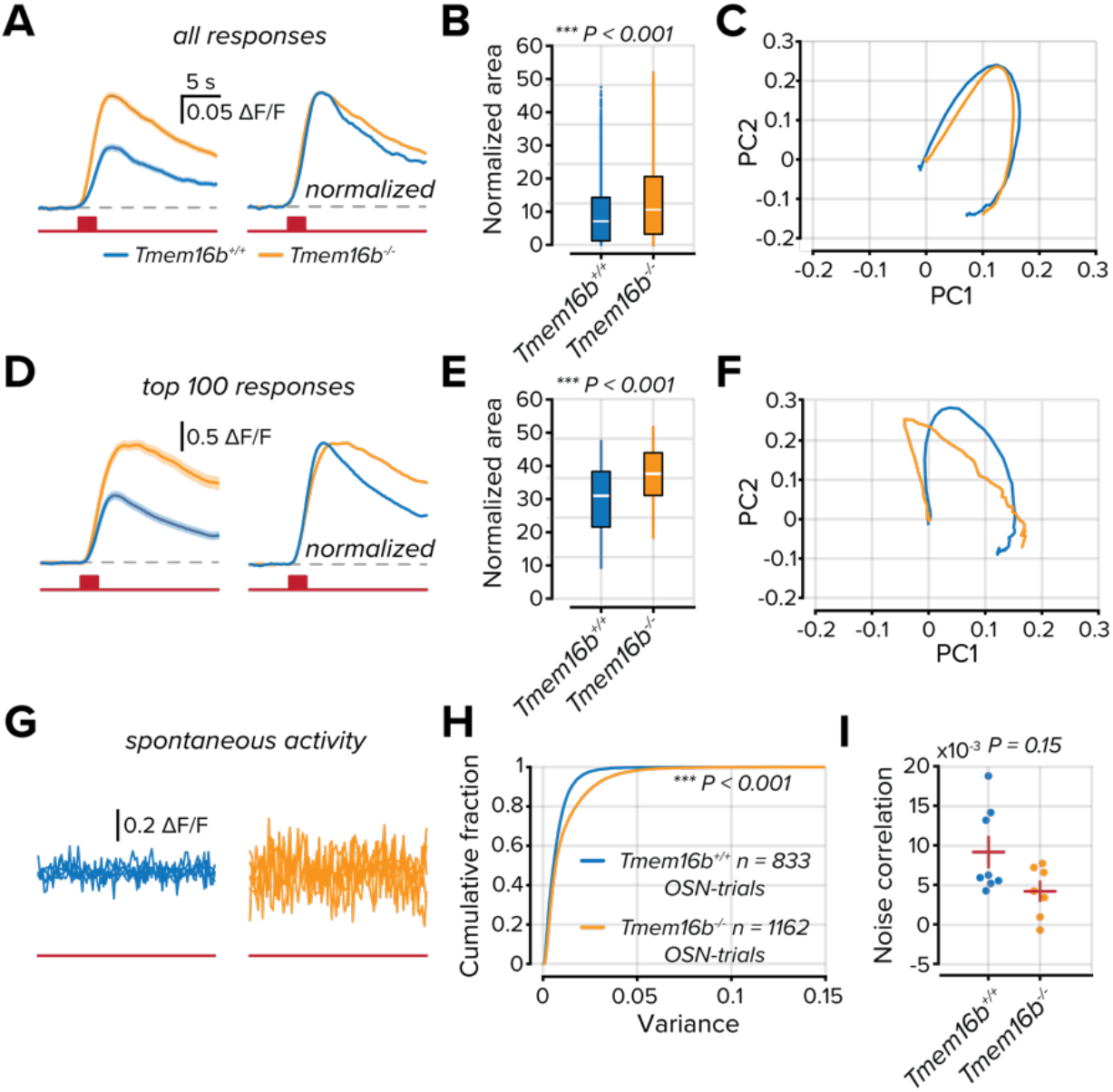
TMEM16B attenuates the temporal dynamics of OSN activation. **A**. Mean responses of all modulated OSN responses. *Right*, peak normalized mean OSN responses. The solid line is the mean response, and the shaded area is the standard error of the mean. The red line below indicates the entire duration of the trial, and the red box corresponds to a 2 s odorant delivery period. **B**. Area measured from the peak normalized OSN responses. The white line represents the median. **C**. Plot of the first two principal components from peak normalized OSN responses. **D-F**. Same as *part A-C* but considering only the 100 largest responses from each group. **G**. Five overlaid traces from separate OSNs from no odorant trials. The red line below indicates the trial duration. **H**. Cumulative distribution of the variance measured in the ΔF/F signals from all OSNs in no odorant trials. **I**. Correlations between OSNs in each imaging field on no odorant trials. The red horizontal line represents the mean, and the vertical line represents the standard error of the mean.

Despite *Tmem16b*^*-/-*^ OSNs having, on average, larger odorant responses, there is a substantial overlap between the activity distributions of the two groups (**Figure 2F**). To evidence the effect on the most responsive units, we compared the temporal profiles of the 100 largest responses from each population (**Figure 4D**). The difference in the decay of the Ca^2+^ transients in *Tmem16b*^*-/-*^ OSNs was markedly greater than in *Tmem16b*^*+/+*^ OSNs (peak normalized area 30.04 ± 0.96; in *Tmem16b*^*+/+*^ OSNs and 37.5 ± 0.75 in *Tmem16b*^*-/-*^ OSNs; *n* = 100; *P* < 0.001; Kolmogorov-Smirnov test; **Figure 4E**). This was also reflected in the PCA trajectories, with apparent differences seen primarily in the first component (**Figure 4F**).

An additional contributor to the density of OSN signaling is their spontaneous activity. To compare spontaneous activity, we collected trials where no odorants were delivered (**Figure 4G**). We measured the variance of the collected ΔF/F signals over the entire trial. OSNs in *Tmem16b*^*-/-*^ mice had greater variance; suggesting a greater degree of spontaneous OSN activity (mean variance = 0.007 ± 0.003; *n* = 833 in *Tmem16b*^*+/+*^ OSNs and 0.008 ± 0.003; *n* = 1168 in *Tmem16b*^*-/-*^ OSNs; *P* < 0.001; Kolmogorov-Smirnov test; **Figure 4H**). As a final component of our analysis, we tested whether spontaneous activity was more correlated in patches of *Tmem16b*^*-/-*^ olfactory epithelium by measuring noise correlations of all OSNs within an imaging field (*n* = 8 *Tmem16b*^*+/+*^ and 7 *Tmem16b*^*-/-*^ imaging fields). We found no difference in the average correlation between groups, indicating that the increased level of spontaneous activity in *Tmem16b*^*-/-*^ OSNs is not uniform and does not arise from a global source.

### *Perceptual switching of odorant valence is shifted in Tmem16b*^*-/-*^ *mice*

Thus far, our findings demonstrate that TMEM16B sparsens sensory input to the olfactory system but limits OSN firing rates and output duration. We, therefore, wondered how *Tmem16b*^*-/-*^ mice would interact with sensory environments of increasing stimulus density. We used an inhibitory avoidance assay with a biphasically-valenced odorant, trimethylamine (TMA). At low concentrations, TMA is attractive to mice; however, as its concentration increases, it becomes strongly aversive (Li et al., 2013).

Mice were allowed to freely explore a two-chamber arena for five minutes, after which volatilized TMA was delivered to one of the chambers for an additional five minutes (*see Methods*; **Figure 5A**). We then compared the time fraction each mouse spent in the odorized and non-odorized chambers at increasing concentrations of TMA. At the lowest two concentrations (0.1% v/v and 1% v/v), there was no difference in place preference between *Tmem16b*^*-/-*^ and *Tmem16b*^*+/+*^ mice (*P* > 0.05; rank sum test; **Figure 5B**,**C**) and both groups displayed neutral or attractive responses to the odorant. However, when the concentration of TMA increased to 10% v/v, we observed a clear difference between groups. *Tmem16b*^*-/-*^ mice had an aversion to the odorized chamber, while *Tmem16b*^*+/+*^ mice were attracted (time fraction in the odorized chamber = 0.60 ± 0.08 in *Tmem16b*^*+/+*^ mice and 0.28 ± 0.04; *n* = 6 mice, both groups; *P* = 0.009; rank sum test). At the highest concentration (100% v/v), both groups equally avoided the odorized chamber (*P* > 0.05; rank sum test).

**Figure 5.**
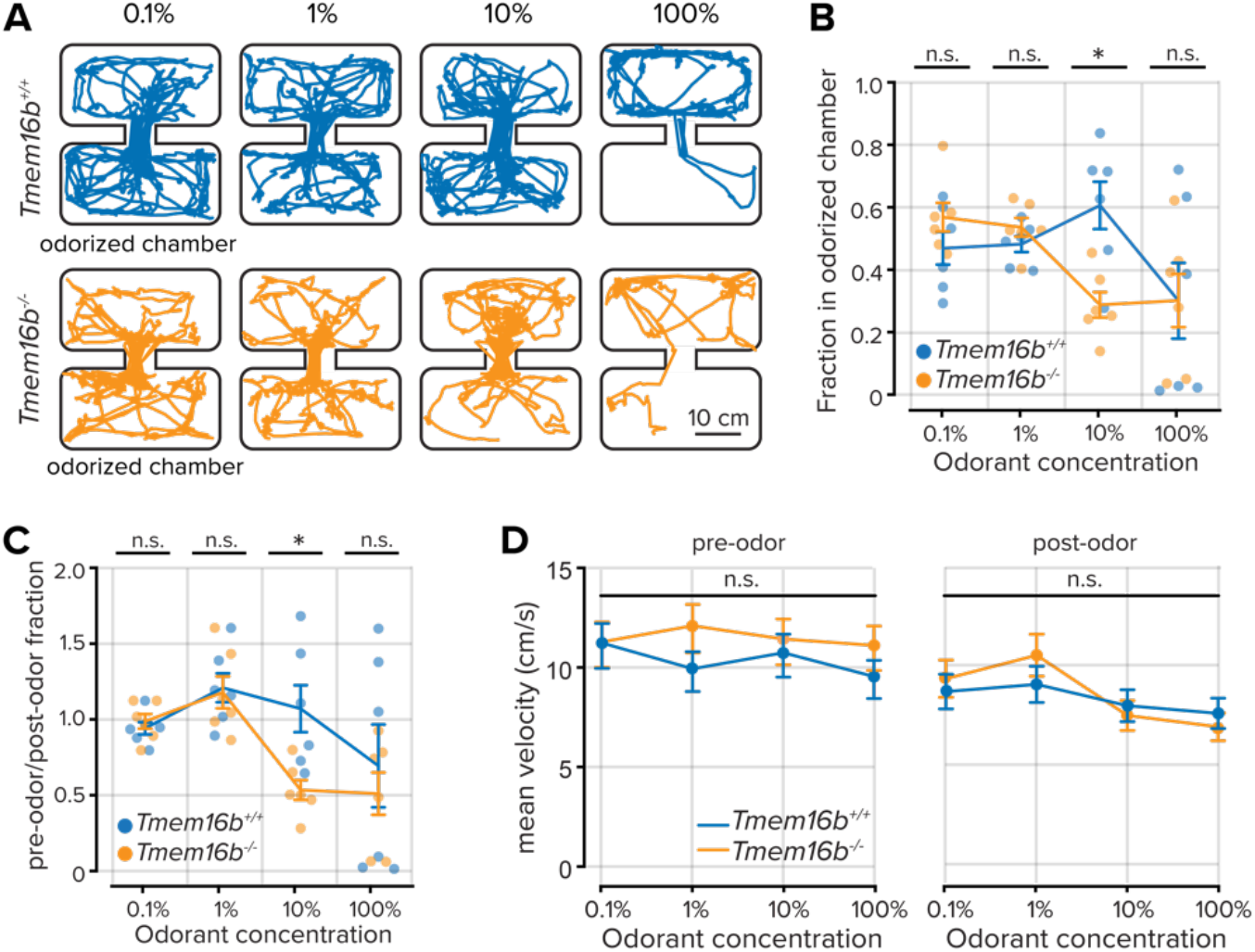
Shifted perceptual thresholds in Tmem16b^*-/-*^ *mice*. **A**. Locomotor paths of example *Tmem16b*^*+/+*^ and *Tmem16b*^*-/-*^ mice at increasing concentrations of the odorant trimethylamine. **B**. Plot of the fraction of time spent in the odorized chamber for each odorant concentration. Lines are the mean data, and error bars represent the standard error of the mean. **C**. Plot of the relative time fraction spent in the odorized chamber before and during odorant delivery for each odorant concentration. Lines are the mean data, and error bars represent the standard error of the mean. **D**. Mean velocity of each animal prior to and following odor onset (*P* > 0.05, Wilcoxon rank-sum test). Lines are the mean data, and error bars represent the standard error of the mean.

TMEM16B is also expressed in the cerebellum and contributes to fine motor control, including gait and locomotion (Zhang et al., 2017). We compared the velocity of both groups before and after odorant delivery to ensure that the increased odorant aversion in *Tmem16b*^*-/-*^ mice was unrelated to locomotor effects. We found no difference between the groups (*P* > 0.05; rank-sum test; **Figure 5D**). Our results show that TMEM16B indeed contributes to olfactory-guided behaviors related to the density of sensory inputs. *Tmem16b*^*-/-*^ mice display an increased aversion at lower concentrations of the biphasically-valenced odorant TMA, thereby indicating that TMEM16B contributes to perceptual thresholds.

### *The efficiency of olfactory-guided navigation is concentration-dependent in Tmem16b*^*-/-*^ *mice*

Our previous results demonstrating increased odorant sensitivity in *Tmem16b*^*-/-*^ mice support the hypothesis that TMEM16B contributes to sparse sensory coding. We next wondered how an increased density of sensory inputs might affect integrated olfactory-guided behaviors. We devised a behavioral paradigm that required mice to navigate within a circular arena toward increasing concentrations of the naturally appetitive odorant peanut oil (Zak et al., 2018). The source of the odorant varied along the perimeter of the arena to prevent animals from learning the source location, while mice innately navigated toward the odorant source under IR illumination.

*Tmem16b*^*+/+*^ mice efficiently navigated to the odorant source regardless of the odorant concentration (**Figure 6A,B**). We hypothesized that odorant-guided navigation efficiency in *Tmem16b*^*-/-*^ mice would decrease as a function of odorant concentration due to an increasing density of OSN activity. However, we found that while there was a relationship between odorant concentration and the latency to find the odorant source, *Tmem16b*^*-/-*^ mice navigated to the source more efficiently as the concentration of the odorant increased (**Figure 6A,B**). At the lowest two concentrations (1% v/v and 10% v/v dilutions), *Tmem16b*^*-/-*^ mice took longer to locate the odorant than *Tmem16b*^*+/+*^ mice (mean latency at 1% peanut oil = 28.98 ± 5.65 seconds in *Tmem16b*^*+/+*^ mice; *n* = 11; and 94.71 ± 16.47 seconds in *Tmem16b*^*-/-*^ mice; *n* = 10; *P* = 0.002; rank sum test. Mean latency at 10% peanut oil = 33.60 ± 7.10 seconds in *Tmem16b*^*+/+*^ mice; *n* = 11; and 79.28 ± 20.28 seconds in *Tmem16b*^*-/-*^ mice; *n* = 10; *P* = 0.028; rank sum test). At higher concentrations (50% v/v and 100% v/v dilutions), there was no difference between groups (*P* > 0.05; rank sum test).

**Figure 6.**
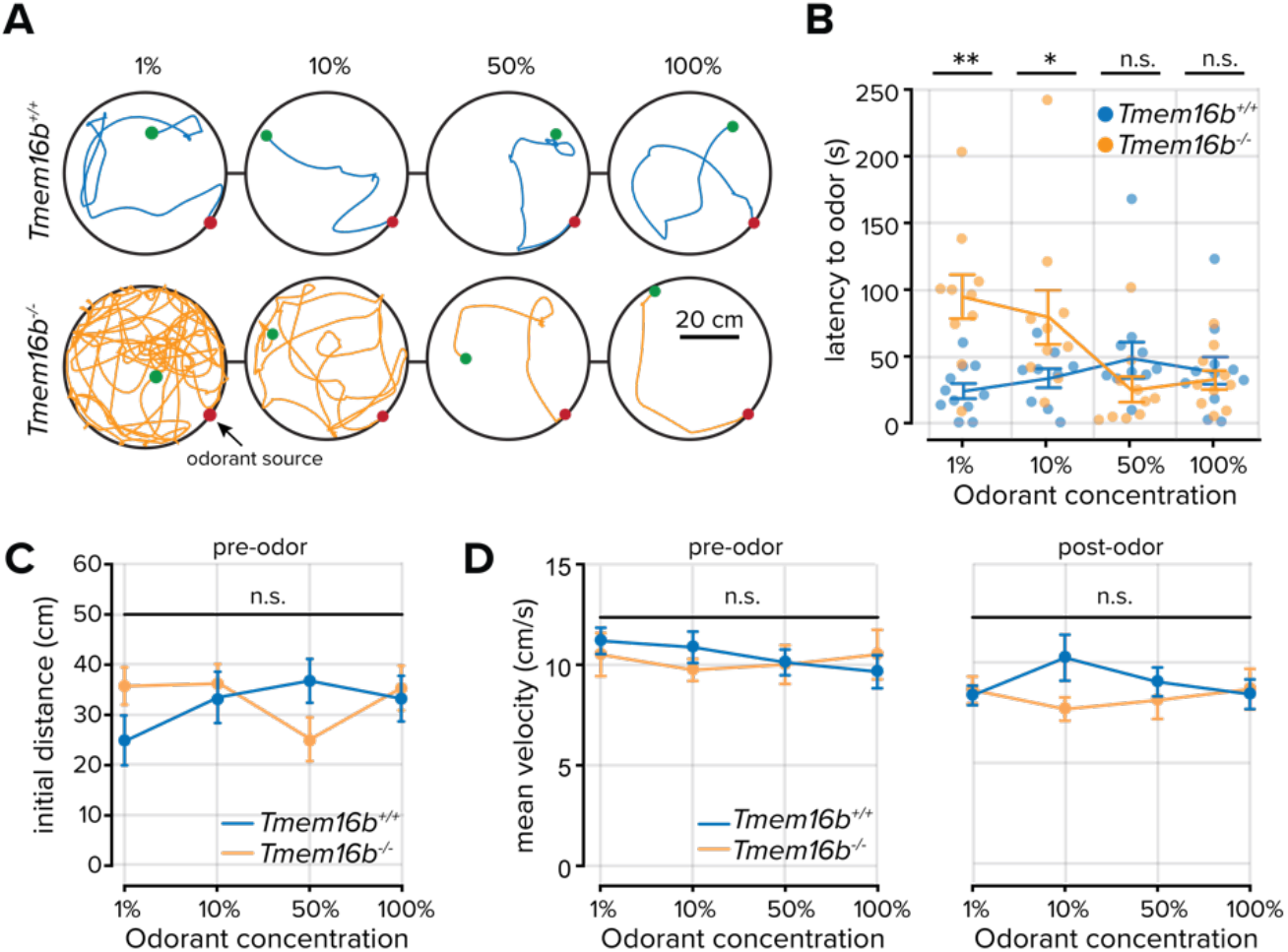
Odorant source localization in Tmem16b^*+/+*^ and *Tmem16b*^*-/-*^ mice. **A**. Locomotor paths of example *Tmem16b*^*+/+*^ and *Tmem16b*^*-/-*^ mice at increasing concentrations of the odorant peanut oil. Green and red dots mark the initial and final position of each animal, respectively. **B**. Plot of the time to locate the odorant for each concentration. Lines are the mean data, and error bars represent the standard error of the mean. **C**. Initial distance from the odorant source (at odor onset) across all animals (*P* > 0.05, Wilcoxon rank-sum test). Lines are the mean data, and error bars represent the standard error of the mean. **D**. Mean velocity of each animal prior to and following odor onset (*P* > 0.05, Wilcoxon rank-sum test) Lines are the mean data, and error bars represent the standard error of the mean.

To account for differences in the mouse position at the opening of the odorant valve, we measured the initial distance from the mouse to the odorant source and found no difference between groups (all comparisons *P* > 0.05; rank-sum test; **Figure 6C**). Again, we compared the velocity of both groups before and after odorant delivery to ensure that the increased latency in locating the odorant in *Tmem16b*^*-/-*^ mice was unrelated to locomotor effects. We found no difference between the groups (all comparisons *P* > 0.05; rank-sum test; **Figure 5D**). Our results demonstrate that TMEM16B indeed contributes to fine-scale olfactory-guided behaviors related to the density of sensory inputs. However, unexpectedly, the navigation efficiency of *Tmem16b*^*-/-*^ mice increased with odorant concentration.

## Discussion

The olfactory epithelium detects volatile odorant molecules, often at low concentrations. The multistep olfactory transduction cascade amplifies transduction currents such that relatively few odorant molecules can exert large electrochemical effects on membrane potentials. In this study, we show that transduction amplification primarily functions to sparsen OSN output rather than boost cellular output and contributes to olfactory-guided behaviors and perception. At the level of individual OSNs within the olfactory epithelium of live mice, we found that TMEM16B reduces the population activity of sensory neurons. The reduction in OSN output increases affective perceptual thresholds, yet simultaneously, allows for efficient integrative behaviors.

### There is a lack of evidence for the dual role of TMEM16B

Despite the increased odorant-evoked activity we observed in *Tmem16b*^*-/-*^ OSNs, these cells had lower lifetime sparseness values, indicating decreased odorant selectivity. What accounts for this paradoxical relationship? One potential explanation is that TMEM16B may function as an amplifier in some situations. For instance, when odorant concentrations are low or receptor-ligand affinities are weak, TMEM16B may amplify transduction currents to levels necessary to generate OSN output without overamplifying and acting as a shunt.

Our imaging experiments used monomolecular odorants at a single dilution (1% v/v); however, our odorant panel covered a range of vapor pressures spanning six odors of magnitude. Therefore, the effective concentration range of our odorant panel spanned a wide range. We found a general relationship between vapor pressure and OSN activity (**Figure 2G**), yet we failed to find evidence at the lowest effective concentration odorants that *Tmem16b*^*-/-*^ OSNs had weaker responses than *Tmem16b*^*+/+*^ OSNs (**Figure 2H**). This is also consistent with another recent report that measured OSN responses across concentrations of a single odorant (Zak et al., 2018). Regardless of odorant concentration, TMEM16B limits OSN output.

A more parsimonious explanation for the reduction in lifetime sparseness is that the threshold for identifying modulated OSN responses is influenced by the higher rate of spontaneous activity we measured in *Tmem16b*^*-/-*^ OSNs (**Figure 4G**). We classified OSNs as being modulated when their odorant responses exceeded three standard deviations of the baseline noise. Our criteria and the higher baseline noise in *Tmem16b*^*-/-*^ OSNs likely mean that weak responses were classified as non-responsive trials, resulting in the appearance of more narrow stimulus tuning. The increased spontaneous activity in *Tmem16b*^*-/-*^ OSNs is also likely to be physiologically relevant, as increased baseline activity contributes to the overall density of information broadcast from the sensory periphery to downstream processing centers.

### Fine-scale olfactory behaviors are affected in Tmem16b^*-/-*^ mice

Consistent with our physiological measurements of OSN activity, our behavioral studies indicate that *Tmem16b*^*-/-*^ mice have an increased aversion to an odorant that changes valence from attractive to aversive by avoiding it at lower concentrations than *Tmem16b*^*+/+*^ mice. Our physiological experiments of OSN odorant responses support this finding. Odorants with the lowest vapor pressures reliably drove stronger peripheral activity in *Tmem16b*^*-/-*^ OSNs, indicating that TMEM16B acts as an amplifier in limited scenarios, at best. TMEM16B sparsens peripheral sensory responses and decreases stimulus sensitivity.

What are the consequences for integrative olfactory-guided behaviors where animals must make decisions based on temporal comparisons of a sensory environment? In a second series of behavioral experiments, we challenged mice to locate the source of peanut oil, an innately appetitive odorant, in a chamber devoid of other sensory cues. Here, we found a dramatic reduction in the ability of *Tmem16b*^*-/-*^ mice to localize the odorant at low concentrations. Interestingly, even at these low concentrations, we still observed behaviors consistent with odorant detection in *Tmem16b*^*-/-*^ mice, including freezing, rearing, and casting (see **Supplemental Video 1**). These observations suggest that *Tmem16b*^*-/-*^ mice can detect the presence of the odorant in the chamber but are unable to perform other computations necessary for localization.

Our study and others provide evidence that TMEM16B constrains the temporal profile of OSN output (Reisert et al., 2024). Discretizing bouts of OSN activity within the respiratory cycle is likely to be essential for making comparisons between sampling events (sniffs; discussed in Fyke et al., 2024). In *Tmem16b*^*-/-*^ mice, odorant samples may be obscured by blending multiple sniffs together and convolving the temporal and concentration gradients found in odorant plumes (Celani et al., 2014; Rigolli et al., 2022). Future studies of respiration and OSN temporal dynamics will be important for understanding how ionic conductances, like those through TMEM16B, contribute to coding features, including phase locking in OSNs.

### Normalization and homeostatic plasticity in downstream circuits

What accounts for the concentration-dependent improvement in odorant localization in *Tmem16b*^*-/-*^ mice? If TMEM16B sparsens OSN population activity and constrains their temporal dynamics, it is perhaps unexpected that localization of an odorant source is most affected at the lowest odorant concentrations. Olfactory information is processed in a multistep hierarchy from the OB to the olfactory cortex. At each processing node, local circuits are positioned to normalize incoming information from the periphery (Aungst et al., 2003; Arevian et al., 2008; Bolding and Franks, 2018; Bolding et al., 2020; Zak and Schoppa, 2022). For instance, granule cells and short axon cells provide targeted inhibition to OB output neurons (Lledo et al., 2008; Burton, 2017), and in the piriform cortex, dense recurrent inhibition implements concentration invariance (Bolding and Franks, 2018). Each of these circuit elements is well-positioned to homeostatically compensate for an increased density of peripheral sensory inputs by tuning inhibitory synaptic weights and densities.

Cell-intrinsic properties of OSNs may also be tuned to account for their increased excitability. OSN axon terminals have a high release probability (Murphy et al., 2004); therefore, the readily releasable pool of synaptic vesicles may be depleted early in a sustained spike train in the absence of TMEM16B. How these circuit and cell-autonomous mechanisms contribute to perception and stimulus evaluation in persistently dense sensory environments has yet to be addressed.

Numerous studies have explored how sensory-depleted environments contribute to circuit development and connectivity (Lorenzon et al., 2015; Galliano et al., 2021; George et al., 2022). The sensory hyperexcitability, evident in *Tmem16b*^*-/-*^ mice, provides a unique model to study dense sensory inputs and their effects on circuit maturation and efficient stimulus coding.

## Methods

### Experimental model and subject details

*Tmem16b*^*+/+*^, *Tmem16b*^*-/-*^, and *OMP-GCaMP3* mice on a C57Bl/6J background of both sexes were used in this study. Sex was not considered in the study design. Mice were acquired from the Jackson Laboratory (Stock #033836) or breeding stocks at Harvard University (OMP-GCaMP3) and maintained within Harvard University’s Biological Research Infrastructure or the Biological Research Laboratory at the University of Illinois Chicago for the duration of the study. All animals were between 20 and 30 g before surgery and singly housed following any surgical procedure. Animals were between three and six months old at the time of the experiments. All mice used in this study were housed in an inverted 12-hour light cycle at 22 ± 1 °C at 30–70% humidity and fed ad libitum.

### Ethics oversight

All experiments were performed in accordance with the guidelines set by the National Institutes of Health and approved by the Institutional Animal Care and Use Committee at Harvard University (protocol 29-20) or the University of Illinois Chicago (protocol 22-011).

### Bone thinning over the olfactory epithelium

Mice were anesthetized with an intraperitoneal injection of ketamine and xylazine (100 and 10 mg/kg, respectively), and the eyes were covered with petroleum jelly to keep them hydrated. Body temperature was maintained at 37 °C by a heating pad. The scalp was shaved and then opened with a scalpel blade. The cranial bones over the olfactory epithelium, anterior to the frontonasal suture, and between the internasal and nasal-maxillary sutures were thinned with a dental drill and scalpel blade until transparent. The thinned area of the skull was then covered with cyanoacrylate adhesive (Loctite 404 Quick Set Adhesive), and a glass coverslip was implanted in the adhesive. The posterior portion of the exposed skull was gently scratched with a blade, and a titanium custom-made head plate was glued (Loctite 404 Quick Set Adhesive) on the scratches. C&B-Metabond dental cement (Parkell, Inc.) was then used to form a well over the thinned section of the skull (Zak, 2022). After surgery, mice were treated with carprofen (6 mg/kg) and buprenorphine SR-Lab (1.0 mg/kg). All animals were allowed to recover for at least three days before imaging experiments were initiated. Throughout the imaging experiments, mice were anesthetized with a ketamine/xylazine mixture to stabilize respiration.

### Multiphoton imaging

A custom-built two-photon microscope was used for *in vivo* imaging. Fluorophores were excited and imaged with a water immersion objective (20X, 0.95 NA, Olympus) at 920 nm using a Ti:Sapphire laser with dispersion compensation (Mai Tai HP, Spectra-Physics). Images were acquired at 16-bit resolution and 4-8 frames/s. The pixel size was 0.6 μm and fields of view were 180 × 180 μm. The point-spread function of the microscope was measured to be 0.51 × 0.48 × 2.12 μm. Image acquisition and scanning were controlled by custom-written software in LabView (National Instruments). Emitted light was routed through two dichroic mirrors (680dcxr, Chroma, and FF555-Di02, Semrock) and collected by photomultiplier tubes (R3896, Hamamatsu) using filters in the 500–550 nm range (FF01–525/50, Semrock).

### Odor stimulation

Monomolecular odorants (Sigma or Penta Manufacturing) were used as stimuli and delivered by custom-built 16-channel olfactometers controlled by custom-written software in LabView (Albeanu et al., 2018; Zak et al., 2024). For all experiments, the initial odorant concentration was 16% (v/v) in mineral oil and further diluted 16 times with air. The airflow to the animal was held constant at 100 mL/min, and odorants were injected into a carrier stream. Odorants were delivered two to seven times each in a trial-based structure. In each trial, a five-second baseline period was followed by a two-second odorant delivery period. The intertrial interval between odorant deliveries ranged between 20-30 seconds.

The odor panel consisted of: 1) Ethyl tiglate 2) Allyl tiglate 3) Hexyl tiglate 4) Methyl tiglate 5) Isopropyl tiglate 6) Citronellyl tiglate 7) Benzyl tiglate 8) Phenylethyl tiglate 9) Ethyl propionate 10) 2-Ethyl hexanal 11) Propyl acetate 12) 4-Allyl anisole 13) Ethyl valerate 14) Citronellal 15) Isobutyl propionate 16) Allyl butyrate 17) Methyl propionate 18) Pentyl acetate 19) Valeric acid 20) (+)Carvone 21) (−)Carvone 22) 2-Methoxypyrazine 23) Isoeugenol 24) Butyl acetate 25) Valeraldehyde 26) Isoamyl acetate 27) Methyl valerate 28) Octanal 29) 2-Hexanone 30) Methyl butyrate 31) 2-Heptanone 32) Acetophenone. Further odorant information is available at ref. (Zak et al., 2020)

### Image processing and data analysis

Images were processed using both custom and available MATLAB (Mathworks) scripts. Motion artifact compensation and denoising were done using NoRMcorre (Pnevmatikakis and Giovannucci, 2017). The CaImAn CNMF pipeline (Giovannucci et al., 2019) was used to select and demix ROIs. ROIs were further filtered by size and shape to remove merged cells. The mean ΔF/F signal in the 5 s following odorant onset was used to measure neural activity in all experiments. To account for changes in respiration and anesthesia depth, correlated variability was corrected (Mathis et al., 2016). Thresholds for classifying responding ROIs were determined from a noise distribution of blank (no odorant) trials from which three standard deviations were used for responses. In each dataset, only ROIs with at least one significant odorant response were included for further analysis. Representational similarity between stimuli was estimated by calculating the Pearson correlation coefficient between population vectors that consisted of all ROIs that satisfied the thresholding criterion.

Sparseness measures were calculated as reported by ref. (Wallace et al., 2017). Lifetime sparseness measures the extent to which a given element responds to different odorant stimuli. Values near one indicate all odorants uniformly activate a given element, and values near zero indicate a high degree of odorant selectivity:

*Equation 1*

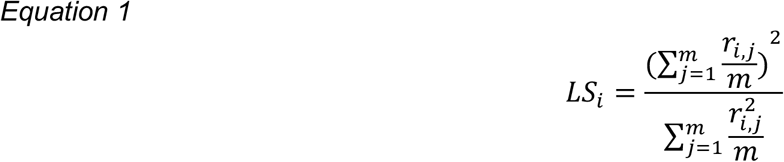

Where: *m* = the number of odorants, *r*_*i,j*_ = the response of a given OSN *i* to odorant *j*.

Population sparseness measures the fraction of or cells that are activated by a given odorant, with values near one indicating uniform activity across all elements and values near zero indicating a lack of activity in most elements:

*Equation 2*

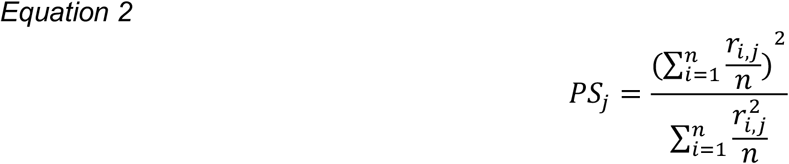

Where: *n* = the number of cells, *r*_*i,j*_ = the response of a given OSN to odorant *j*.

All statistical comparisons for imaging experiments were made as described in the text for each figure and values are given as mean ± standard error of the mean.

### Inhibitory avoidance behavioral assessment

The behavior apparatus consisted of two semi-transparent 18 cm × 30 cm × 12 cm chambers connected via a short 6 cm diameter tube. The apparatus is housed inside a sound and light-proof outer chamber illuminated with LED lights. Both chambers of the apparatus had two valve-controlled air inlets, allowing air and odor to flow at a constant velocity of < 100 mL/min. After 5 minutes of baseline exploration, the air inlet to one side of the chamber was switched to an odor inlet using a manual valve. A clear acrylic cover was placed on top of the apparatus to contain mice and odorant in the chambers. Trimethylamine (CAS: 75-50-3, Sigma Aldrich) was serially diluted in mineral oil to four concentrations (0.1%, 1%, 10%, and 100% v/v). Each animal was tested for 10 minutes in a randomized order and once per concentration. The chambers were thoroughly vacuumed for at least 5 minutes and cleaned with 70% ethanol after each trial to clear the chambers of any odor accumulation and presence of social cues. Images were acquired at ten Hz using a USB camera (DFK 42BUC03, The Imaging Source) with an accessory camera lens (IP/CCTV 13VM308ASIRII, Tamron) and processed using custom-written MATLAB scripts to measure the location of the midpoint of the mouse body, which was then used to calculate velocity. All statistical comparisons for behavior experiments were made with the Wilcoxon rank-sum test and values are given as mean +/-standard error of the mean.

### Open Field Behavior

The arena consisted of a 56 cm diameter circular inner chamber with four air inlets equally spaced around its circumference. The circular inner chamber was housed in a light- and sound-proof outer chamber and illuminated using infrared LEDs. Throughout each experiment, airflow was maintained at a constant velocity for each inlet. After 10 minutes of baseline exploration, air to one of the inlets was redirected through an odorized chamber while ensuring no change in its velocity. A vacuum was located at the center of the arena, and its flow matched the sum of all air inlets to prevent the accumulation of odor in the arena. Peanut oil was diluted in mineral oil as in imaging experiments. Each animal was only tested once per odorant concentration, and the order in which they were tested was randomized. After each trial, the arena was thoroughly cleaned with 70% ethanol to eliminate the presence of social cues. Images were acquired at eight Hz using a USB camera (Grasshopper3, Point Grey Imaging) and custom-written LabVIEW software. Images were processed using custom MATLAB routines to measure location and velocity. Animals with an initial position < 10 cm from the odor source were excluded from our analysis. All statistical comparisons for behavior experiments were made with the Wilcoxon rank-sum test and values are given as mean +/-standard error of the mean.

## Data and Code Availability

The data and code supporting this study’s findings will be deposited in a GitHub database upon publication.

## Acknowledgments

We thank Venkatesh Murthy for sharing equipment and resources. Anna Vlasits, Angles Salles, and all members of the Zak Laboratory for helpful discussions. This work was supported by NIH Grant R00 DC017754 to JDZ. VK was supported by NIH Grant T32 DK128782.

## Author Contributions

JDZ, KCB, and EH designed the experiments. JDZ, KCB, EH, ZF, LV, and VK collected the data. JDZ, KCB, and EH analyzed the data. JDZ wrote and edited the manuscript with input from all authors.

