## Supplementary figures and images for "Amplification of olfactory transduction currents implements sparse stimulus encoding"

### Supplemental Video 1

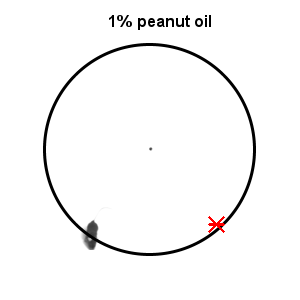
